# BMP antagonist CHRDL2 enhances the cancer stem-cell phenotype and increases chemotherapy resistance in Colorectal Cancer

**DOI:** 10.1101/2024.01.23.576664

**Authors:** Eloise Clarkson, Annabelle Lewis

## Abstract

BMP antagonists have been increasingly linked to the development of Colorectal cancer (CRC). BMP signalling operates in opposition to the WNT signalling pathway, which sustains stem-cell maintenance and self-renewal of the normal intestinal epithelium. Reduced BMP and elevated WNT signalling lead to expansion of the stem-cell compartment and the hyperproliferation of epithelial cells, a defining characteristic of CRC. Chordin-like-2 (CHRDL2) is a secreted BMP antagonist, with overexpression linked to poor prognosis and variants in the gene shown to be associated with an elevated CRC risk. Despite this the functional role of CHRDL2 in CRC is unknown.

In this study, we explored the impact of CHRDL2 overexpression on CRC cells to investigate whether CHRDL2’s inhibition of BMP signalling intensifies WNT signalling and enhances the cancer stem-cell phenotype and response to treatment. Our research approach combines 2D cancer cell lines engineered to inducibly overexpress CHRDL2 and 3D organoid models treated with extrinsic CHRDL2, complemented by RNA sequencing analysis.

CHRDL2 was found to enhance the survival of CRC cells during chemotherapy and irradiation treatment due to activation of DNA damage response pathways. Organoids treated with secreted CHRDL2 exhibited elevated levels of stem-cell markers and reduced differentiation, as evidenced by diminished villi budding. RNA-seq analysis revealed that CHRDL2 increased the expression of stem-cell markers, WNT signalling and other well-established cancer-associated pathways.

These findings collectively suggest that CHRDL2 overexpression could affect response to CRC therapy by enhancing DNA repair and the stem-cell potential of cancer cells, and its role as a biomarker should be further explored.

**Graphical abstract:** 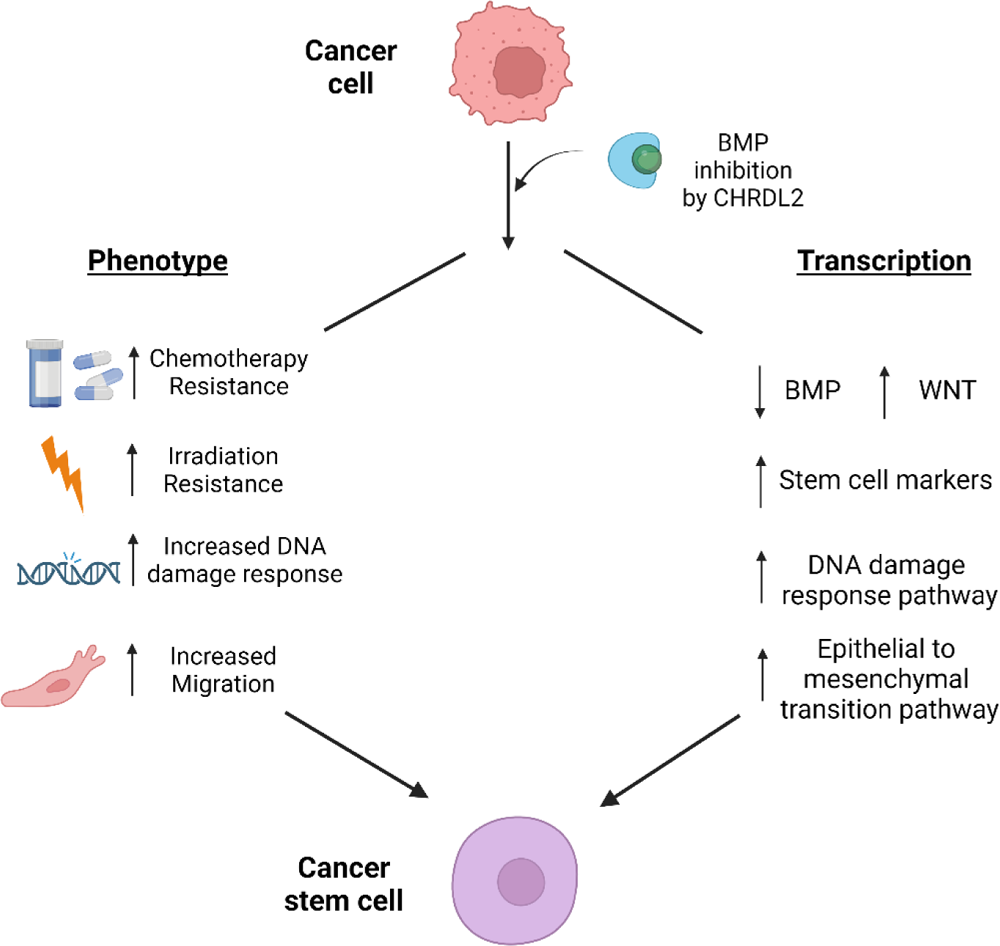

## Introduction

Colorectal cancer (CRC) ranks as the third most prevalent cancer globally, with over 1 million reported cases in 2020 by GLOBOCAN [1]. Originating from mutations within intestinal epithelial cells, CRC leads to the formation of polyps, adenocarcinomas and eventually metastatic cancer. Although many signalling pathways are disrupted in CRC, the WNT/β-catenin signalling pathway is the most commonly affected, overactive in nearly all CRC cases [2].

WNT signalling is the fundamental pathway regulating intestinal stem-cell (ISC) proliferation and fate. WNT activation is localized at the base of the intestinal crypt, where it controls ISC fate and renewal, a vital component of intestinal maintenance [3]. Cryptal ISCs are organised hierarchically, with rapidly proliferating ISCs residing at the bottom of crypts and more slowly proliferating or regenerative ISCs slightly displaced from the base of the crypt at the +4 position [4,5]. A third more mature subset of rapidly proliferating cells migrate further up the crypt into the transit amplifying (TA) zone. ISCs are identified by the expression of the LGR5+ marker, and slow-cycling cells in the +4 position have previously been identified by the presence of the BMI1+ marker^5^, as well as HPOX and TERT, but with conflicting data about the specific markers and role for these cells [6].

WNT signalling transduces by sequestering the β-catenin destruction complex, raising intracellular β-catenin levels, and activating stem-cell and oncogenic pathways in stem-cell crypts [7,8]. Disruption of WNT signalling, eliminates progenitor phenotypes within the crypts, and results in crypt loss [9]. Despite the need for WNT signalling to maintain the intestinal lining and crypt formation, sustained or elevated WNT-signalling can cause hyperproliferation and oncogenic transformation in ISCs [10]. WNT signalling has been shown to work in a counter gradient to BMP signalling, which is found in the intestinal villi and promotes cellular differentiation and maturation [10–12]. These gradients of BMP and WNT signalling are a major controlling factor in crypt-villi architecture and intestinal homeostasis.

In contrast to WNT signalling, BMP signalling is localized in the differentiated compartment of the crypt/villi and aids in cellular differentiation, proliferation, and migration [13]. BMPs have been shown to have paradoxical effects in cancer, with specific ligands acting to inhibit and promote tumorigenesis in different tissues and contexts [[14–17]. BMPs belong the TGF-β superfamily and bind to a complex of transmembrane serine threonine kinase receptors I and II (BMPRs I and II) [18]. This initiates phosphorylation of the type I receptor by the type II receptors, triggering phosphorylation of a receptor-associated SMAD that subsequently complexes with SMAD4 and translocates to the nucleus to regulate gene transcription [19]. While epithelial and mesenchymal cells express BMPs and their receptors, BMP antagonists are primarily found in the mesenchyme. In the intestine they are largely expressed by intestinal cryptal myofibroblasts and smooth muscle cells. These antagonists block BMP signalling in the stem-cell compartment, maintaining high levels of WNT signalling and therefore the stem-cells [11].

BMP antagonists can bind directly to BMPs or their receptors [20]. Some well-studied BMP antagonists include Noggin, which has been implicated in promoting skin and breast cancer tumorigenesis, and the Gremlins (GREM1 and 2), with repression of GREM1 shown to inhibit tumour cell proliferation. The Chordin family of proteins have also been implicated in CRC, including Chordin, Chordin-like 1 (CHRDL1) and Chordin-like 2 (CHRDL2) [21–24]. One of the best studied BMP antagonists, Noggin, has previously been shown to inhibit BMP signalling in a mouse model, resulting in the formation of numerous ectopic crypts perpendicular to the crypt-villus axis [12]. Similarly, overexpression of GREM1 in Hereditary mixed polyposis syndrome (HMPS) leads to the persistence or reacquisition of stem-cell properties in LGR5-negative cells outside the stem-cell niche. Ectopic crypts, enhanced proliferation and intestinal neoplasia [25]. Together, this suggests that abolition of BMP signalling through its antagonists leads to the formation of stem-like qualities in intestinal epithelial cells, leading to oncogenic transformation.

CHRDL2 is a BMP antagonist which prevents BMP ligands, most likely BMP2 and BMP4, from interacting with their cognate cell-surface receptors [22,26]. CHRDL2 has been shown to bind directly to BMPs, preventing signalling though phospho SMAD1/5. Furthermore, CHRDL2 inhibits the effects of BMP signalling on proliferation inhibition and apoptosis [27]. CHRDL2 mRNA upregulation has been observed in colon, breast, liver, and prostate cancer [27,28] and high levels predict poor prognosis, and correlate with increased tumour size and later TNM stages [27]. CHRDL2 has been highlighted as a potential circulating protein biomarker for CRC, in which genetically predicted higher levels of CHRDL2 were associated with an increased risk of CRC [29]. CHRDL2’s precise functional role in these cancers is not always clear but it has been shown to increase cellular proliferation, migration, and invasion in osteosarcoma cell lines by regulation of the PI3k/AKT pathways through binding to BMP9 [30]. However, the role of BMP signalling in cancer, and therefore the effect of BMP inhibition by CHRDL2 in cancer remains poorly characterised.

In this study we used CRC cell lines engineered to stably overexpress CHRDL2 in an inducible manner, to investigate the cellular and transcriptional pathways activated by CHRDL2 expression and BMP inhibition. We have shown that CHRDL2 has measurable effects on cell proliferation and significantly changes the response to DNA damaging chemotherapy. To gain deeper insights into CHRDL2’s role in stem-cell maintenance and differentiation, we cultivated 3D intestinal organoids supplemented with secreted forms of CHRDL2. Collectively, our findings suggest that CHRDL2 modulates stem-cell pathways in CRC, potentially impacting the response to common chemotherapeutic interventions.

## Results

### CHRDL2 overexpression inhibits cell proliferation and decreases clonogenicity

To determine the effects of CHRDL2 on colorectal cancer cells, we transduced four extensively characterised CRC cell lines with a virally packaged doxycycline-inducible overexpression system for full-length *CHRDL2* cDNA. Colorectal adenocarcinoma cell lines were deliberately chosen to encompass a range of *CHRDL2* expression levels and genetic mutations: CACO2 and LS180 (moderate CHRDL2), COLO320 and RKO (very low). Doxycycline was given in 3 concentrations to cell lines: 0.1 μg/ml (CHRDL2), 1 μg/ml (CHRDL2+) or 10 μg/ml (CHRDL2++) (Figure 1A) to induce expression. qPCR and Western blotting confirmed overexpression of CHRDL2 at the RNA and protein level respectively (Figure 1B, C, D). Conditioned media from CHRDL2-overepxressing cell lines was also collected, and secreted CHRDL2 protein was found to be present in the media. BMP antagonism was shown through assessing levels of phosphorylated SMAD 1/5, which occurred in LS180, RKO and COLO320 (Figure 1G, Supplementary Figure 1).

**Figure 1:**
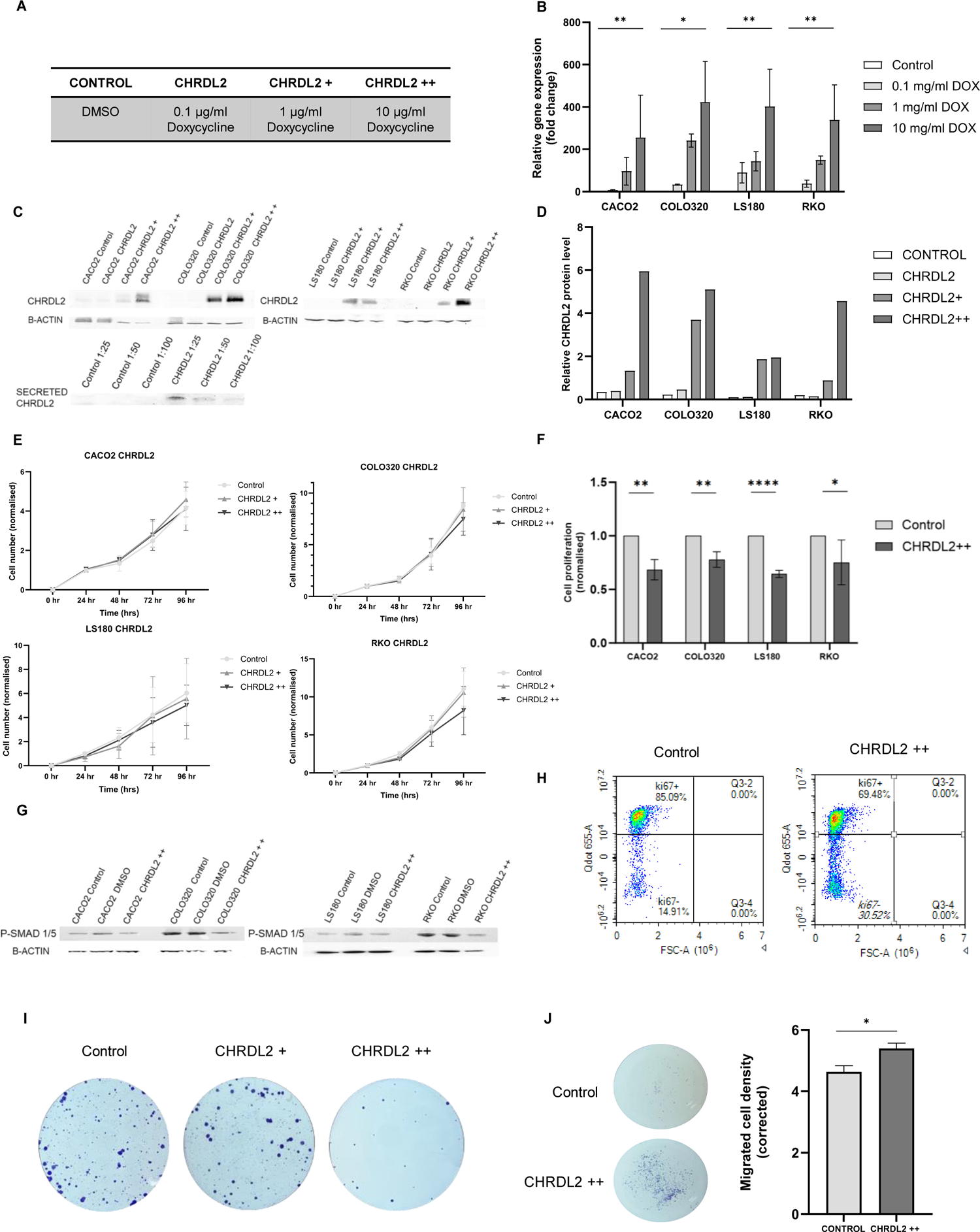
Inducible CHRDL2 overexpression in CRC cell lines alters proliferation and migration A) Table of doxycycline treatment used, B) qPCR of mRNA levels of CHRDL2 expressed as fold change in 4 experimental cell lines. Cell lines were grown with doxycycline at: 0.1 μg/ml, 1 μg/ml or 10 μg/ml to induce expression. RKO DMSO-10μg/ml p<0.01, COLO320 DMSO-10μg/ml p<0.05, CACO2 DMSO-10μg/ml p<0.01, LS180 DMSO-10μg/ml p<0.01. N=3, T-test. C) Western blotting of corresponding protein levels of CHRDL2 in cell lines with lentiviral overexpression, and secreted CHRDL2 present in cell culture media. D) Quantification of CHRDL2 protein levels as measure by western blot using Image J software. E) MTT assay of cellular proliferation of CHRDL2 cell lines. In, COLO320 and RKO cells lines, small but significant decreases in CHRDL2 expression were shown. Two-way RM ANOVA P<0.01, P<0.036 respectively F) Cellular proliferation analysis on cells grown in low glucose conditions, given 10 μg/ml doxycycline overexpression of CHRDL2. In CACO2, COLO320, LS180 and RKO cells lines, proliferation was significantly decreased with CHRDL2 expression. T-test; P<0.01, P<0.01, P<0.001 and P<0.01 respectively. G) Western blotting of SMAD1/5 phosphorylation in cell lines overexpressing CHRDL2. H) Flow cytometry analysis of COLO320 cells given CHRDL2++ overexpression using Ki67 antibody staining as a marker of proliferating cells. Proportion of Ki67 cells decreased with CHRDL2++ overexpression, I) Crystal violet staining of colonies of RKO cells treated with 1 μg/ml and 10 μg/ml doxycycline to induce CHRDL2 expression. J) Crystal violet staining of migrated COLO320 cells with 10 μg/ml doxycycline to induce CHRDL2 expression. Quantification on ImageJ shows significant increase in number of migrating cells with CHRD2++, P<0.0449. Error bars given as ± SEM.

Increased proliferation is a hallmark of cancer cells. Therefore, we measured the effects of CHRDL2 overexpression on cellular proliferation in our cell lines. As seen in figure 1E, cell growth was slightly reduced during overexpression of CHRDL2 in COLO320 and RKO (P<0.01, P<0.036,). Intriguingly, this effect was enhanced when cells were cultured under low glucose conditions; high levels of CHRDL2 (++) expression reduced proliferation in all our cell lines: CACO2 (p<0.01), LS180 (p<0.001), RKO (p<0.05), and COLO320 (p<0.01) (Fig. 1F).

As seen in Figure 1H, CHRDL2 overexpression decreased the proportion of proliferating cells, marked by Ki67+ (Supplementary Figure 2A, p<0.05). In Ki67+ populations, CHRDL2 increased the number of S phase cells and lowered the proportion in G2 phase, possibly reflecting a decreased rate of proliferation (Supplementary Figure 2B, p<0.01). Investigation of the colony forming competency (clonogenicity) of CHRDL2 overexpressing cells revealed that clonogenic potential was reduced (Figure 1I). This was found in all four tested cell lines (Supplementary Figure 2C, p<0.01). Overall, contrary to our hypothesis, CHRDL2 overexpression appears to reduce proliferation and colony formation. However, our observation that the effects of high CHRDL2 are enhanced by reduced levels of glucose, suggests a more complex phenotype. This could perhaps be due to a preference for aerobic glycolysis, the Warburg effect. This is a known characteristic of cancer stem-cells and may suggest that CHRDL2 causes an increase in some stem-cell characteristics to accompany a decrease in proliferation rate [31].

Migratory ability is a further measure of stem-cell competency, so to measure this in cells with CHRDL2 overexpression, COLO320 cells were seeded in transwell inserts. The number of cells migrating through a porous membrane to the lower chamber was quantified. As seen in Figure 1J, CHRDL2 overexpression significantly increased the number of migrated cells (P<0.0449,), suggesting increased migratory ability, a hallmark of cancer stem-cells.

### CHRDL2 increases resistance to common chemotherapy

Another characteristic of cancer stem-cells is resistance to chemotherapy [32]. In light of this, our study aimed to evaluate the response of our experimental cell lines to the three most common chemotherapy agents employed in the treatment of CRC.

We treated CHRDL2++ cells with chemotherapy drugs, 5-Fluorouracil (5FU), irinotecan and oxaliplatin, and assessed cellular response via MTS assay. Figure 2A, shows the reduction in cell number with increasing chemotherapy concentration (μM). Control cell lines were plotted together with cell lines with CHRDL2 overexpression, and the half maximal inhibitory concentration (IC50) values were calculated (Figure 2A, B, C and Supplementary Figure 3). CHRDL2 overexpression significantly increased resistance to chemotherapy in all cell lines (P<0.01) as shown by elevated IC50 values (Figure 2B). CHRDL2 overexpression resulted in an approximate twofold increase in IC50 values compared to control cells, (P<0.001). Average increases in IC50 values during CHRDL2 overexpression for each drug and cell lines can be observed in Figure 2C. The greatest increase in survival (exhibited by ratios of IC50s) was seen in COLO320 cells treated with oxaliplatin, which had a 3.6 fold increase.

**Figure 2:**
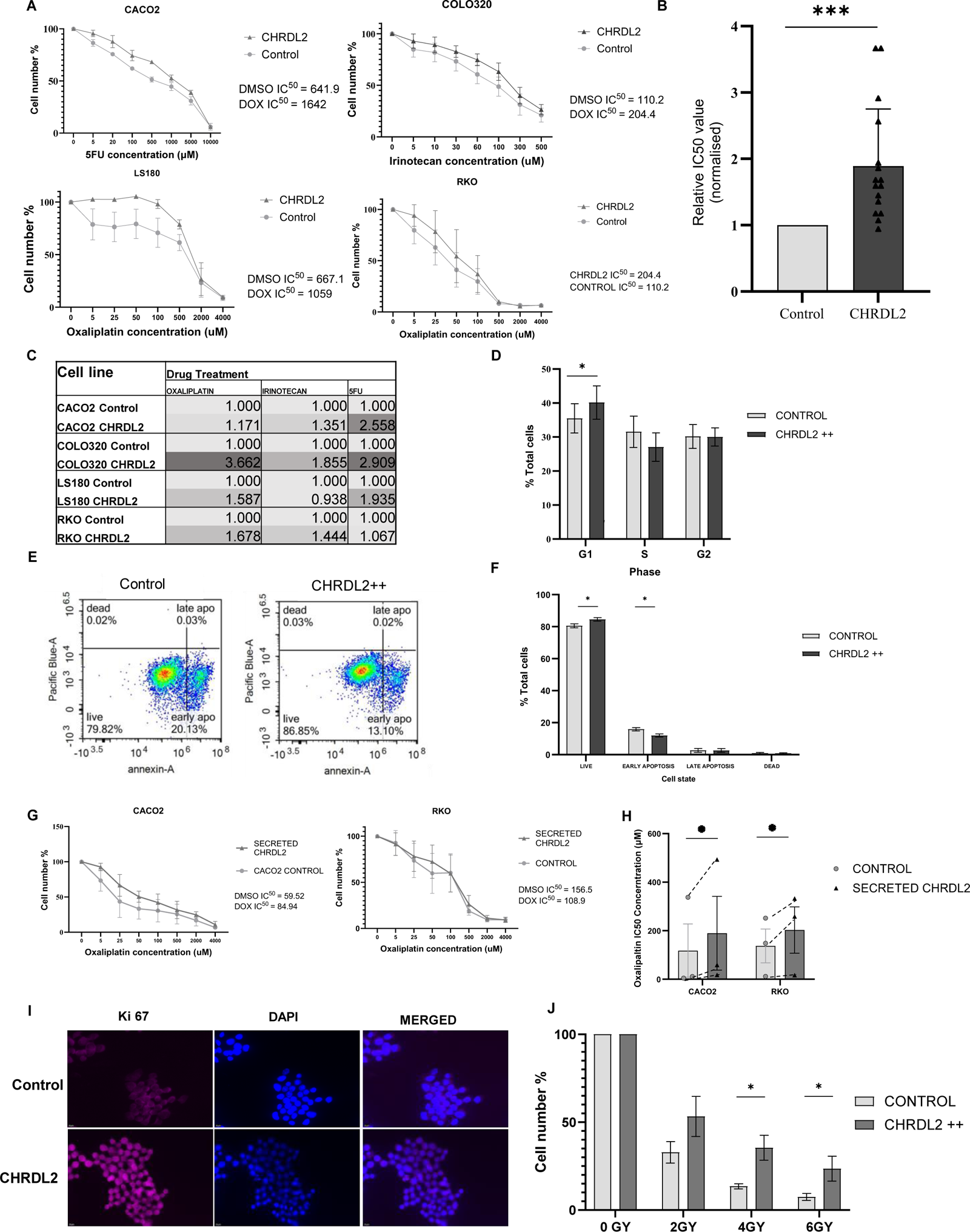
CHRDL2 overexpression increases resistance to common CRC chemotherapies. A) Drug dose response curves using CACO2 cells and 5FU, COLO320 cells and Irinotecan, and LS180 and RKO cells and Oxaliplatin N=3. Two-way ANOVA was used to find differences between curves, P<0.0068, P<0.0001, P<0.0006, P<0.005 respectively. B) Average difference in IC50 valued across all cell lines and 3 chemotherapy drugs. P< 0.005. C) Table of ratio differences in IC50 values between CHRDL2++ cells and control for each chemotherapy drug and cell-line. N=3. D) Flow cytometry analysis of COLO320 cells treated with Oxaliplatin and CHRDL2 Overexpression. CHRDL2 increased the number of cells in G1 phase. (students T-test P<0.003). E) Flow cytometry analysis of COLO320 cells treated with Oxaliplatin and CHRDL2 overexpression stained with Annexin-5 as a marker of apoptosis. F) Quantification of cell percentages of live, apoptotic and dead cells in COLO320 cells treated with oxaliplatin and given CHRDL2 overexpression. CHRDL2 increased the % of live cells (students T-test P<0.03) and decreased the % of early apoptotic cells (P<0.023). N=3. G) Drug dose response curves of CACO2 and RKO cells with CHRDL2 conditioned media and Oxaliplatin N=3. Two-way ANOVA was used to find differences between curves, P<0.005, P<0.0001 H) Average IC50 values for chemotherapy drug Oxaliplatin, on cell lines CACO2 and RKO with CHRDL2 conditioned media. CHRDL2 against control p< 0.0305. N=3. I) Immunofluorescence staining of Ki67 on COLO320 cells treated with 5 μM oxaliplatin. J) Cell count after irradiation of RKO cells overexpressing CHRDL2++ N=3. T-test 4GY: P<0.038, 6GY: P<0.0241. Error bars given as ± SEM.

Flow cytometry of COLO320 cells treated with oxaliplatin (Figure 2D) showed a clear increase in the number of cells in S phase (in both control and CHRDL2++ cells) compared to the untreated cells shown in Figure 1J. This is probably due to stalling of replication forks due to DNA damage exerted by chemotherapy, and activation of the S/G2 checkpoint. Interestingly, cells with CHRDL2 overexpression displayed a smaller increase in cells stalled in S Phase compared to controls, and therefore showed a greater proportion of present in G1 phase. This is the opposite to that of untreated cells in Figure 1J, where CHRDL2 increased the number of cells in S Phase possible due to slower cell division. Further flow cytometry analysis following chemotherapy treatment revealed CHRDL2 overexpression decreased the number of cells that had entered early apoptosis (P<0.05) (Fig 2E and F) demonstrating that CHRDL2 overexpressing cells have the ability to evade apoptosis.

Secreted CHRDL2 from conditioned media was also used on our parental non-CHRDL2 expressing cell lines to assess paracrine signalling. Again, secreted CHRDL2 increases cellular survival during chemotherapy in the same manner as our intracellular CHRDL2 overexpression system, P<0.005 with elevated IC50 values (Fig 2G and H.)

To further analyse the survival capabilities of CHRDL2 overexpressing cells during chemotherapy, Ki67 staining was performed. As seen in figure 2I, CHRDL2 overexpression significantly increased the number of proliferating cells during chemotherapy treatment, shown by an upregulation in Ki67 (P<0.0064, Supplementary 4B). This is supported by flow cytometry analysis (P<0.0055, Supplementary 4A).

Resistance to irradiation, along with chemotherapy resistance, is also attributed to more aggressive cancers that evade treatment. ICSs have been shown to have increased resistance to irradiation, therefore we sought to measure the effects of CHRDL2 overexpression on cell survival during X-ray irradiation. Cells were treated with 0 GY, 2 GY, 4 GY or 6 GY X-ray irradiation and cell viability was assessed. As seen in figure 2J, CHRDL2 overexpression increased cell survival at 4GY and 6GY radiation (P<0.03, P<0.02,).

### CHRDL2 overexpression decreases DNA damage during chemotherapy treatment

Next, we investigated the mechanism through which CHRDL2 promotes cell survival during chemotherapy treatment. The chemotherapy agent Oxaliplatin is known to cause DNA intra-strand cross-linking, resulting in double strand breaks (DSBs), cell-cycle arrest and apoptosis [33]. Therefore, quantification of DSBs in cells treated with Oxaliplatin was performed on COLO320 CHRDL2++ cells using immunofluorescence staining of γH2AX and Ku70. Quantification of DNA repair proteins ATM and RAD21 was also performed, to assess whether CHRDL2 protects cells from DNA damage through upregulation of DNA repair pathways.

Figure 3A images show DSBs in cells treated with a low dose of Oxaliplatin (approx. IC25) after 24, 48, and 72 hrs post treatment. Quantification using Image J demonstrated that CHRDL2 overexpressing cells had significantly fewer γH2AX foci compared to the control at each time point (Figure 3B P<0.01, t-test). This difference was notably increased at 72 hours compared to 24 hours. This suggests that CHRDL2 does not necessarily protect cells from DNA damage but rather acts to accelerate the repair of DNA damage when compared to control cells. Similarly, after 72hrs there was a decrease in the presence of Ku70, which binds to DSBs to facilitate non-homologous end-joining (NHEJ), in COLO320 CHRDL2 cells (P<0.0057, Figure 3C and supplementary 4C). This further confirms a reduction in DNA damage through CHRDL2 upregulation.

**Figure 3:**
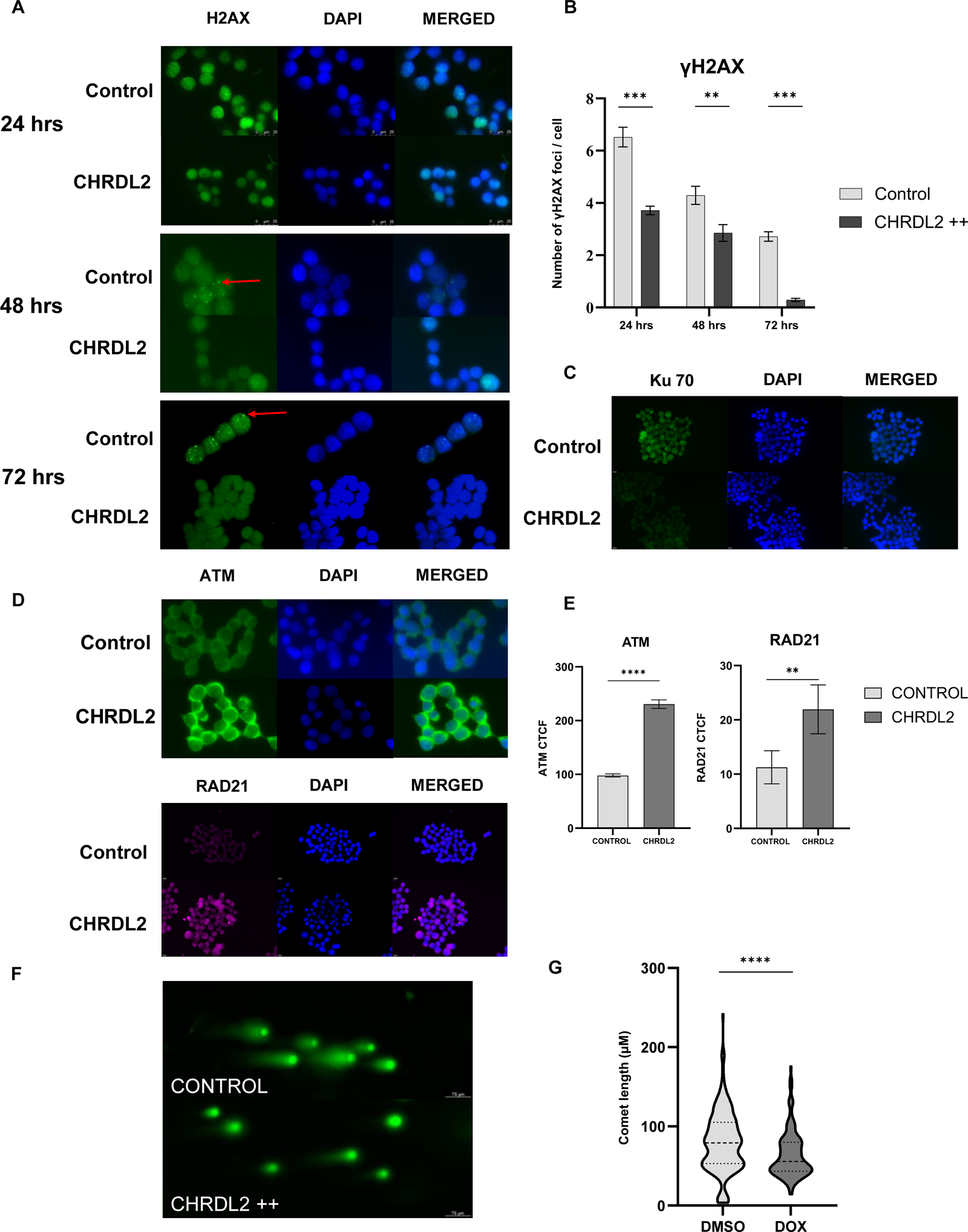
CHRDL2 overexpression decreases DNA damage during chemotherapy treatment and enhances expression of DNA repair pathways. A) Representative immunofluorescence of γH2AX on COLO320 cells treated with 5 μM oxaliplatin at 24, 48 and 72 hrs. Foci indicated by red arrows B) Quantification of γH2AX foci in COLO320 cells overexpressing CHRDL2 treated with 5 μM oxaliplatin at 24, 48, and 72 hrs. Cells were treated with DMSO control reagent, or Doxycycline to induce CHRDL2++ overexpression. 24hs P< 0.0001, 48hrs P<0.01, 72hrs P<0.0001. N=3. T-test C) Immunofluorescence staining of Ku 70 on COLO320 cells treated with 5 μM oxaliplatin. D) Immunofluorescence staining of ATM and RAD21 on COLO320 cells treated with 5 μM oxaliplatin. E) Quantification of ATM and RAD21 staining on COLO320 cells. Immunofluorescence given as Corrected Total Cell Fluorescence (CTCF). Cells were treated with DMSO control reagent, or Doxycycline to induce CHRDL2++ overexpression. P<0.0001 and P<0.0023 respectively. N=3 T-test. F) Comet assay of RKO cells treated with IC50 Oxaliplatin. Cells were then treated with CHRDL2 ++ overexpression or a control. G) Quantification of Comet assay, T-test P<0.0001. N=3. Error bars given as ± SEM. Quantification carried out using Image J.

This is supported by Figure 3D and E, in which we observed significantly (P<0.0001) increased ATM, and RAD21 (P<0.0023) in CHRDL2 overexpressing cells compared to the control, suggesting upregulation of DSB damage response pathways in which ATM serves as a master transducer. γH2AX is also known to accumulate during cellular senescence. However, since we found no difference in P53 expression, (a marker of senescence, supplementary Figure 5) in our CHRDL2 overexpressing cells it is likely that upregulation of DNA damage pathways in CHRDL2++ cells protects against DNA damage by chemotherapy.

We have further demonstrated the ability of CHRDL2 overexpression to reduce DNA damage during chemotherapy by alkaline comet assay, as observed in figure 3F and G. Cells were treated in the same manner with IC25 Oxaliplatin. We observed cells with CHRDL2 overexpression had shorter “tails” to their comets, showing less fragmented or damaged DNA. Quantification using ImageJ confirmed this, with CHRDL2++ cells having significantly decreased tail length compared to control cells (P<0.0001).

### CHRDL2 decreases organoid budding and increases stem-cell markers

CRC cell lines exhibit a multitude of abnormalities, characterized by numerous mutations, heightened WNT signalling, and impaired DNA repair mechanisms. Consequently, we also explored the impact of CHRDL2 overexpression on normal ISCs within an organoid model, aiming to shed light on the role of CHRDL2 in tumour initiation.

To replicate the stem-cell niche and overexpression of CHRDL2 paracrine signalling, which typically originates from mesenchymal cells, we established murine intestinal organoids, providing a three-dimensional platform for modelling the effects of CHRDL2. These organoids were exposed to secreted forms of CHRDL2 in the form of conditioned media obtained from our cell lines. As a control, conditioned media from parental (non-CHRDL2 expressing) cells undergoing doxycycline treatment was also collected.

Figure 4 A showcases the distinctive morphological features of intestinal organoids, characterized by a villi-bud-like configuration, with the outer epithelial layer forming distinct protrusions and invaginations. Prior investigations have examined the gene expression profiles of intestinal organoids, revealing that epithelial cells within the “buds” of the organoids exhibit crypt-like expression patterns, while the evaginations display villus-like expression [34].

**Figure 4:**
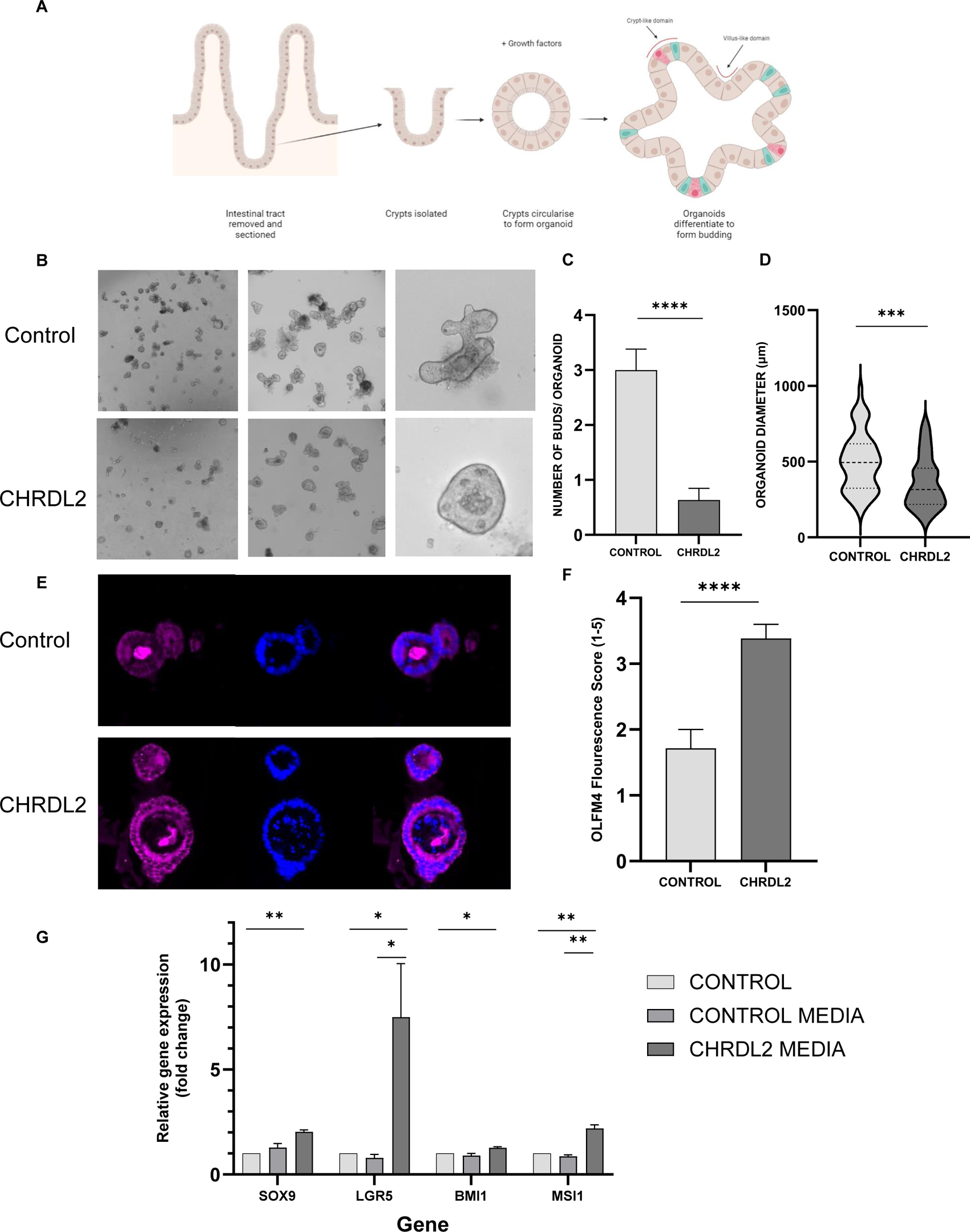
Secreted CHRDL2 decreases murine small intestinal organoid differentiation and increases stem-cell marker expression. A) Image of intestinal organoid diagram depicting organoid structure. B) Image of murine-derived organoids treated with conditioned media containing secreted forms of CHRDL2 compared to conditioned media from control cells with no CHRDL2 overexpression. C) Quantification of buds per organoid in CHRDL2 treated murine organoids compared to a control. T-test P<0.0001. D) Quantification of average organoid diameter in CHRDL2 treated murine organoids compared to a control. T-test P<0.001. E) Immunofluorescence staining of OLFM4 on murine organoids treated with secreted CHRDL2 compared to a control after 1 week. F) Quantification of immunofluorescence scoring of OLFM4 on murine organoids treated with secreted CHRDL2 compared to a control. T-test P<0.0001. G) qPCR of stem-cell markers from CHRDL2 treated murine organoids compared to a control. T-test SOX9. Students T-test P<0.0014, LGR5 P<0.04, P<0.043, BMI1 P<0.0113, MSI1 P<0.0067, P<0.009. Error bars given as ± SEM.

As seen in figure 4 B, upon the addition of extrinsic CHRDL2 to organoids, a noticeable reduction in the number of differentiated buds was observed (P< 0.001) (Figure 4B and C). Organoids developed smaller and more circular characteristics, suggesting slower growth similar to our observations in cell-lines (P<0.001) (Figure 4B and D). This is supported by the presence of Olfactome-din-4 (OLFM4), a marker for LGR5+ stem-cells [35], which was increased in CHRDL2 treated organics (P<0.0010) (Figure 4, E, F). Moreover, as illustrated in Figure 4G, we observed a significant increase compared to the controls (P<0.05) in the expression of stem-cell markers LGR5 (indicating crypt CBCs) and BMI1 (slow-cycling crypt stem-cells), and also SOX9 and MSI1 . These findings collectively suggest that exposing intestinal organoids to CHRDL2 diminishes differentiation and enhances stem-cell numbers.

### CHRDL2 enhances cancer stem-cell pathways

To elucidate pathways in which CHRDL2 overexpression acts, RNAseq analysis on CACO2 cells given CHRDL2+ and CHRDL2++ treatment was performed with DMSO treated cells as a baseline. Differential expression analysis was carried out on RNAseq data (Figure 5A). 76 and 145 differentially expressed genes were identified in the CHRDL2 + and CHRDL2++ groups respectivvely, (P<0.05,. Figure 5B). From this we selected the 21 genes that were diffentially expressed in both CHRDL2+ and CHRDL2++ vs control cells for downstream analysis (Figure 5C). qPCR was used to confirm expression changes in biological replicates in both CACO2 and COLO320 cells (Figure 5D).

**Figure 5:**
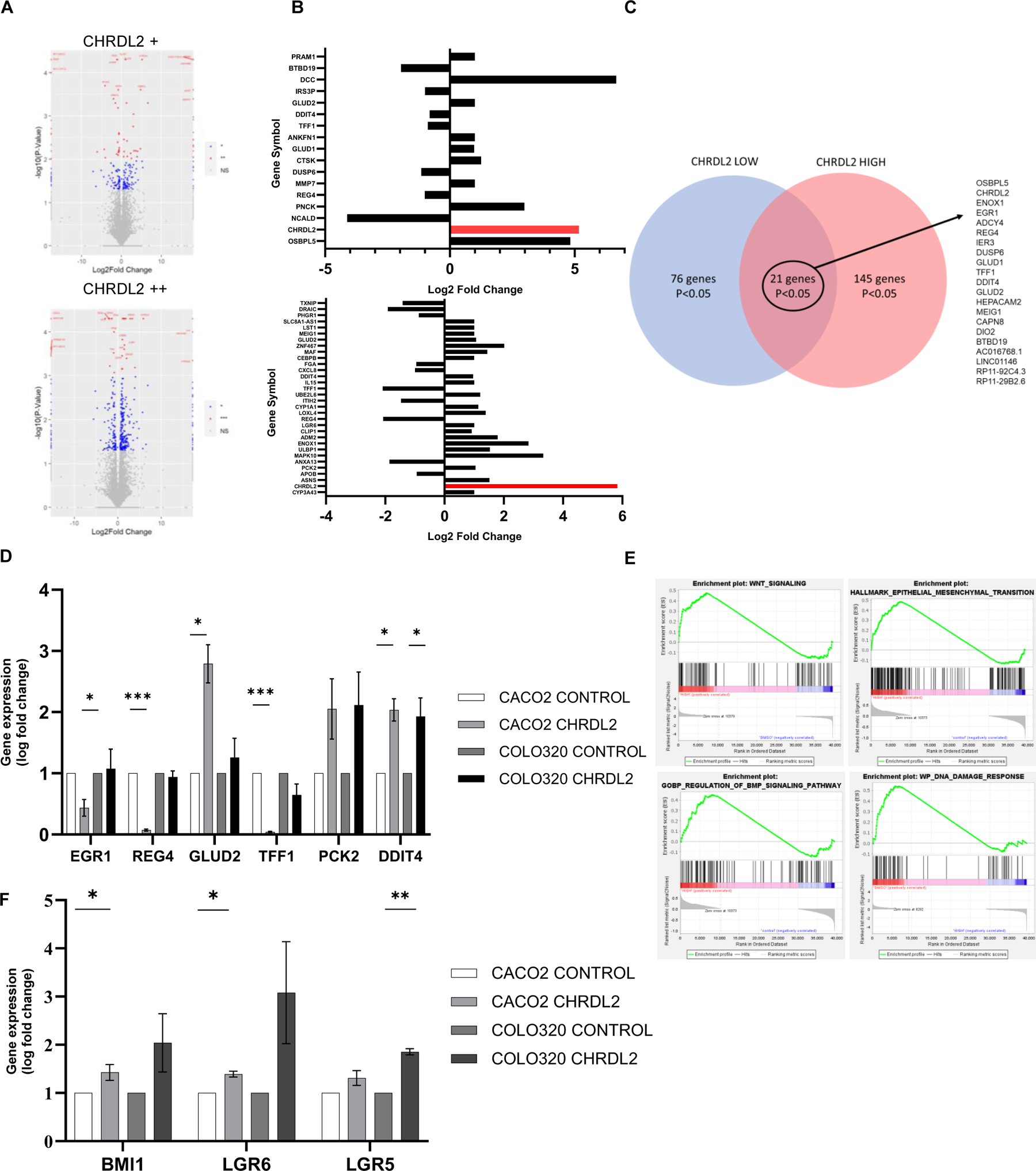
RNAseq analysis demonstrates that CHRDL2 expression enhances cancer stem-cell and other cancer hallmark pathways. RNAseq analysis of CACO2 cells treated with CHRDL2+ or CHRDL2++ compared to a control. N=3 A) Volcano plot of differentially expressed genes from cells with CHRDL2 overexpression (CHRDL2 + and CHRDL2 ++) from RNAseq analysis. Expressed as Log2 Fold change against control. B) Bar-plot of significantly differentially expressed genes in CHRDL2+ and CHRDL2++ cells. Genes included pass the threshold of P<0.001 for CHRDL2+ and P<0.001 for CHRDL2++. C) Intersect of highly differentially expressed genes in both the CHRDL2 + and CHRDL2++ treated groups P<0.001. 21 genes were differentially expressed in both groups. D) qPCR validation of differentially expressed genes from RNAseq data in CACO2 and COLO320 cells N=3.E) GSEA plots of differentially expressed pathways in CACO2 cells treated with CHRDL2++. F) qPCR validation of stem-cell markers in CACO2 and COLO320 cells N=3. CACO2 BMI1 P<0.05, CACO2 LGR6 P<0.05, COLO320 LGR5 P<0.01. N=3. Error bars given as ± SEM.

Interestingly, Trefoil factor 1 (*TFF1*) known to inhibit proliferation, migration and invasion was downregulated in the RNAseq data and in qPCR. Glutamate Dehydrogenase 2 (*GLUD2*), a glycolysis related gene, Phosphoenolpyruvate Carboxykinase 2 (*PCK2*), involved in mitochondrial respiration and elevated in tumours, and DNA damage inducible transcript 4 (*DDIT4*), associated with advanced CRC, were all upregulated in both RNAseq and qPCR data. Stem-cell markers *LGR5* and *LGR6*, as well as B lymphoma Mo-MLV insertion region 1 homolog (*BMI1*) were also upregulated in our qPCR data, with *LGR6*, a WNT transducer, upregulated in RNAseq data in the CHRDL2++ treatment (Figure 5F).

Gene-set-enrichment analysis (GSEA) on the entire RNAseq dataset, revealed upregulation of the hallmark WNT signalling pathway (P<0.001) and BMP regulation (P<0.05). This increase in WNT signalling and decrease in BMP signalling (Figure 5E) verifies CHRDL2’s role as a BMP antagonist in colon cancer cells. The MYC signalling pathway and LEF1 signalling, which are downstream transducers of WNT signalling, were also upregulated. GSEA revealed upregulation of the cancer hallmark pathways, epithelial to mesenchymal transition (EMT) (P<0.001), and angiogenesis, which are frequently upregulated in metastatic colorectal cancers (Supplementary Figure 6). DNA repair pathways, were also significantly upregulated including key DSB repair genes BRCA1, RAD51 and RAD52, supporting our findings with respect to chemo/radiotherapy resistance (P<0.05). There was also significant upregulation of RAF and MTOR signalling, which are often modulated during cancer progression. Furthermore, cell cycle-related genes upregulated by Rb knockout were also upregulated by CHRDL2, suggesting an increase in cell-cycle protein signalling (Supplementary Figure 6). We noted that, BMI1 pathways were also highlighted by GSEA (Supplementary Figure 6) a further stem-cell defining pathway and correlating with our Q-PCR data.

## Discussion

The role of BMP signalling in cancer is well studied but often paradoxical; with BMP signalling shown to be necessary to prevent cancer-associated WNT signalling and ISCs from exiting intestinal-crypts [12,25]. Conversely it can also play a role in promoting tumorigenesis [16]. It is clear, however, that BMP antagonists play an important functional role in regulating BMP signalling and therefore could be a key biomarker in cancer progression [20]. In this study we have confirmed that CHRDL2 represses BMP signalling in CRC cells, leading to elevated WNT signalling, and causes changes in cell growth, response to chemotherapy and stem-cell characteristics.

Previously, CHRDL2 has been shown to bind to BMP2 and 9 to block BMP mediated SMAD1/5 phosphorylation signalling [27,30]. Patient survival studies have shown CHRDL2 overexpression predicts poor prognosis in CRC patients, and mRNA expression is elevated in patient tumour tissues compared to a control. Variation near CHRDL2 has also been implicated as a cause of increased CRC risk in genome-wide-association (GWAS) studies [36] [37]. Knockdown studies of CHRDL2 have been shown to inhibit proliferation and migration in CRC, gastric cancer, and osteosarcoma, and overexpression promotes cellular proliferation, migration and clonogenicity [27,30]. Since other secreted BMP antagonists, Noggin and GREM1, have also been shown to enhance tumorigenesis and modulate intestinal cell stemness, we suggest that CHRDL2 acts in a similar manner and that this is via the BMP pathway [21,23,25].

Here we have used Doxycycline-inducible models to overexpress CHRDL2 at a variety of levels to investigate the transcriptional and behavioural effects of this gene. Despite previous reporting’s of CHRDL2 increasing proliferation, we found a small decrease in proliferation in our CRC cell lines. When we reduced glucose, CHRDL2 overexpression significantly reduced proliferation and also clonogenicity, a measure of both proliferation and colony formation. Some cancer stem-cells (CSCs), including populations in some CRC cell lines, undergo substantial metabolic reprogramming to become glycolytic (the Warburg effect) and are therefore more reliant on glucose [38]. Our observed sensitivity to glucose reduction could be related to enhanced stem-cell properties of cancer cells with CHRDL2 overexpression. It should be noted that glucose-depletion dramatically alters CSC gene expression and cellular behaviour, compared to non-stem cancer cells, which allows them to reduce reactive oxygen species and survive for longer periods of time [39–41].

We also observed enhanced cell migration through a porous membrane of CHRDL2 overexpressing cells. CSCs have long been shown to harbour increased migratory potential, supporting our findings of increased stem like qualities during CHRDL2 overexpression. This is reflected in our RNAseq data, which shows enrichment for the EMT pathway, which relies on enhanced migratory and invasion properties of cancer cells.

Next, we identified increased resistance to the three most common forms of chemotherapy used to treat CRC, again reinforcing the propensity for survival during CHRDL2 overexpression in our CRC cell model. Standard of care for all but the early stages of CRC relies on the use of chemotherapy in combination with surgical procedures. 5-Fluouracil (5FU) is currently the cornerstone of chemotherapy treatment used to treat CRC [42,43] and used in combination with either Oxaliplatin, a diamino cyclohexane platinum compound that forms DNA adducts (Known as FOLFOX) or Irinotecan, a topoisomerase I inhibitor (known as FOLFIRI) [44]. In each cell line we tested, there was an increase in survival during chemotherapy treatment, regardless of mechanism of action of the three chemotherapy agents. This was confirmed through flow cytometry analysis that showed CHRDL2 overexpression reduced the number of cells that entered apoptosis, and also increased the number of proliferating cells remaining after treatment. Additionally, we have shown that CHRDL2 overexpression increases cell survival during irradiation treatment, which can be used concurrently with chemotherapy in the treatment of CRC.

We observed upregulated ATM and RAD21 signalling during chemotherapy of CHRDL2 overexpressing cells indicating hyperactive DNA damage response pathways. This is likely to be a factor in the accelerated clearing of DSBs as marked by the significantly faster reduction in γH2AX foci and Ku70. DNA damage repair genes were also shown to be upregulated by our RNAseq data, as shown by GSEA, although these cells had not been exposed to any chemotherapy and therefore we would expect only baseline DNA repair activity. Exactly which repair pathways are upregulated in response to CHRDL2 cells undergoing chemotherapy, and whether these are error prone or accurate, is an important question that remains to be addressed. In general, enhanced DNA damage response activation is also a hallmark of CSCs, which has been shown extensively to aid CSCs survival following conventional treatments, allowing the return of cancer in patients and worsened prognosis [45–47].

Through comprehensive RNAseq analysis and qPCR validation, we have shown that CHRDL2 overexpression increased WNT signalling, and expression of stem-cell markers, including LGR5, BMI1, LGR6, and SOX9 in 2D and 3D models. This data collectively suggests that CHRDL2 enhances stem-cell capacity through increased WNT signalling, and therefore intensifies a stem-cell like phenotype.

However, CHRDL2 does not enhance proliferation and clonogenicity, suggesting that it could preferentially support cancer stem-cells that are slower cycling as opposed to the hyperproliferative cancer stem-cells. Within the intestinal crypt, normal stem-cells are arranged in a hierarchy, with rapidly proliferating stem-cells or crypt base columnar cells (CBCs) at the base of the crypt. A separate population of stem-cells lie the +4 position, which appear to cycle more slowly [4,5]. Although the data are inconsistent, some studies propose that these slow-cycling stem-cells at the +4 position are marked by BMI1, which was upregulated in our RNAseq and qPCR data, raising the possibility that CHRDL2 enhances this slow-cycling stem-cell phenotype. Slow-cycling CSCs have also been shown to be radiation resistant, similar to our CHRDL2 overexpressing cells [48,49]. There is some conflicting evidence for the role of these slow-cycling stem-cells with a number of publications proposing that they are key for regeneration of the intestine after injury [5,50]. However more recent studies show that LGR5+ CBCs are also able to fulfil this role, or suggest that the two populations support each other to facilitate tissue repair [6].

Our organoid models treated with secreted CHRDL2 further confirmed our increased stem-like hypothesis. When mouse organoids were treated with CHRDL2, they showed a significant reduction in differentiated bud formation, creating a smaller, spherical stem-like phenotype compared to controls. Immunofluorescence staining of stem-cell marker OLFM4 showed upregulation in organoids treated with CHRDL2. Furthermore, stem-cell markers *LGR5*, *MSI1*, *BMI1* and *SOX9* were increased following CHRDL2 treatment showing that inhibition of extracellular BMP signalling can directly increase stemness in normal intestinal cells. In a recent study another BMP antagonist, GREM1, is upregulated in the intestinal stroma in response to injury, resulting in reprogramming/dedifferentiation in intestinal epithelial cells to drive repair [51]. Thus, inhibition of BMP signalling through CHRDL2 or other antagonists may indeed force cancer cells in the colon into a more stem-like state, and while this may not increase proliferation rate it could increase longevity and survival of these cells during treatment with DNA damaging therapies.

Through RNAseq analysis we have also shown differential expression of other cancer biomarkers by CHRDL2, such as *EGR1*, *REG4* and *TFF1*, which have been shown to regulate proliferation, migration, and metastasis. Furthermore, CHRDL2 was found to differentially impact several key cancer pathways, including the EMT pathway, MYC, MTOR, PI3/AKT and RAF. For example, DDIT4 is a regulator of MTOR, and was upregulated by CHRDL2, and EGR1 which acts through PI3K/AKT was downregulated by CHRDL2. Indeed, CHRDL2 has previously been shown to act via PI3K/AKT in osteosarcoma [30]. These data provide new avenues of research into the mechanism that CHRDL2, and potentially other BMP antagonists, may exert their effects. Unravelling the pathways modulated by CHRDL2 and other BMP antagonists will undoubtedly drive future investigations in cancer research.

Our findings suggest that CHRDL2 should be further explored as a potential biomarker for increased chemotherapy resistance in CRC. This would necessitate a deeper understanding of the mechanism of resistance and also accurate determination of the pattern of overexpression of CHRDL2 during tumorigenesis and therapy. In summary, our data strongly suggest that CHRDL2, by inhibiting BMP signalling and augmenting WNT signalling, promotes stem-cell properties in cancer cells, thus contributing to cancer progression and potentially therapeutic resistance.

## Materials & Methods

### Cell culture and maintenance

Immortalised human colorectal adenocarcinoma cell lines CACO2, COLO320, LS180, and RKO (acquired from ATCC) were maintained in Gibco Dulbecco’s Modified Eagle Medium (DMEM) (Sigma-Aldrich) supplemented with 10% foetal bovine serum (FBS) (Sigma-Aldrich), and 1% penicillin streptomycin (Sigma-Aldrich). Cells were grown in an humified atmosphere at 37◦c with 5% CO2. Subculturing was performed every 72 hours to maintain a cell confluency of < 80%.

### Generation and validation of CHRDL2 overexpressing cell lines

CHRDL2 full length cDNA (Genocopoeia GC-H1938) was cloned into pCW57.1 (Addgene #41393) using Gateway technology (Invitrogen, Thermo Fisher, US), followed by validation by Sanger sequencing and restriction digest. The vector was then transfected in HEK293 cells along with viral packaging vectors (2^nd^ generation system – pCMV-dR8.2 and pCMV-VSV-G) using Lipofectamine 2000 (Invitrogen, US). Virus containing media was collected, sterilized and titre measured (Go-Stix, Takara). The cell lines CACO2, COLO320, LS180, and RKO were transduced, and cells with integrated pCW57.1-CHRDL2 were selected with puromycin. To confirm overexpression, doxycycline was added at (0.1 μg/ml, 1 μg/ml (CHRDL2 +) or 10 μg/ml (CHRDL2 ++) RNA was extracted (RNeasy, QIAGEN) and quantified by real-time reverse transcriptase polymerase chain reaction (qPCR) using TaqMan technology (Hs00248808_m1) according to the manufacturers protocol (Applied biosystems). Each assay was repeated in triplicate.

### Western blot

For intracellular protein detection, cells were lysed by resuspension in RIPA buffer. For secreted protein expression, cells given doxycycline expression at 10 mg/ml were incubated for 72 hours. Media was collected and concentrated through Amicon® Ultra centrifugal filters (Merck, UK) with pore size of 30 kDa. 30 μg of protein were loaded per sample. Protein samples were separated via 4-12% sodium dodecyl sulfate polyacrylamide gel electrophoresis under denaturing conditions, and then transferred onto the nitrocellulose membrane (Millipore, UK) under 20 V. Membranes were blocked with 5% milk for 1hr at room temperature. Membranes were then incubated with primary antibody in TBST-5% BSA overnight at 4◦C. Membranes were then washed with TBST. Secondary antibody was added for 1 hr at room temperature. Membranes were imaged through incubation with Enhanced chemiluminescence (ECL). The ratio of optical density of the bands was measured by a gel image analysis system (Bio-Rad) and normalized to B-actin as a loading control.

**Antibodies used**:

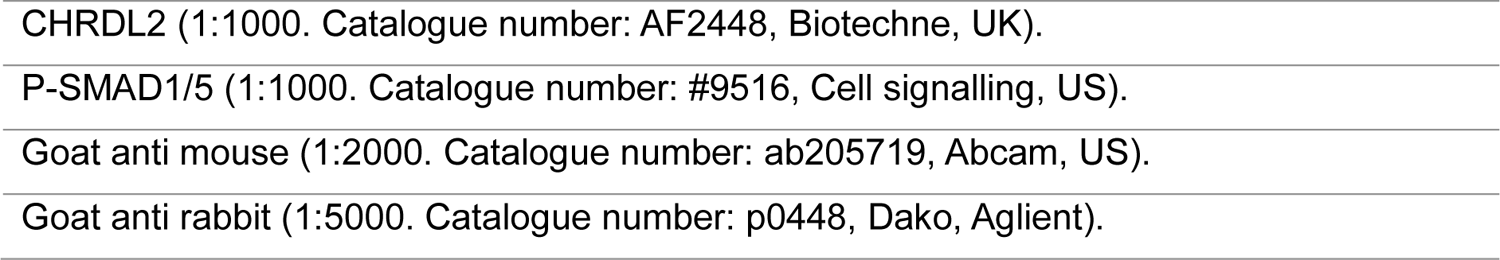

### Cell proliferation assay

To assess cellular proliferation during CHRDL2 overexpression, cells were plated at a density of 5x10^3^ cells per well in 96 well plate with 8 replicates per condition: DMSO control, CHRDL2+ or CHRDL2++ treatments. Cellular proliferation was assessed via MTS assay (CellTiter 96® Promega) at 24, 48 and 72 hrs. For low glucose proliferation, cells were plated in the same manner using glucose deficient medium, and cell number was assayed by MTS after 72hrs. Results were analyzed using Graphpad Prism.

### Flow cytometry

COLO320 cells were plated in six-well plates at a density of 1×10^5^ cells/well under standard media conditions, supplemented with DMSO, or CHRDL2++ treatments. For chemotherapy flow cytometry analysis, cell were treated with 25 μM Oxaliplatin. Cells were grown for 48hrs before harvesting by trypsinization, and washed once with cold PBS. To investigate cell-cycle progression, cells were resuspended in Hoeschst 33342 (62249, Thermo Scientific) and incubated at 37°C for 2hrs with slight agitation. Finally, samples were resuspended in PBS, and flow cytometry analysis was performed using ACEA Novocyte system (Agilent, US). The percentages of cells in phases of the cell cycle were analysed through Novocyte software. To investigate apoptosis, cells were stained with ZombieAqua (423101, Biolegend, US) and annexin-V antibody (V13242, Themo scientific) for 30 mins each at room temperature. Subsequently, cells were washed once with PBS and analysed by flow cytometry using manufacturers software.

### Colony formation assay

To assess the ability of single cells to generate colonies and cell survival ability, a clonogenic assay was performed. Cells were plated at 100 cells per well of a 6 well plate. Cells were treated with doxycycline treatment as before in varying concentrations of 10 μg/ml, 1 μg/ml, and 0.1 μg/ml or DMSO. Plates were incubated for 2 weeks until visible colonies were formed. Every 72 hours doxycycline and DMSO treatments were refreshed. After 2 weeks, cells were fixed with >98% methanol at -20◦C and stained with crystal violet stain (0.5%, in 20% methanol). Colonies were counted through ImageJ.

### Drug dose response assay

To assess the ability of cell lines to withstand treatment from commercial chemotherapy drugs, a drug dose response curve was performed. Chemotherapy drugs used were: 5-Fluorouracil (5FU), Irinotecan, and Oxaliplatin (Sigma Merck). A serial dilution was performed in standard cell-culture media to give a range of concentrations (5FU 0-10000 μM. Oxaliplatin 0-4000 μM, Irinotecan 0-500 μM). Cells from culture were seeded on a 96 well plate at a density of 2 x 10^5^ in 100 µl standard media with 10 mg/ml doxycycline and incubated for 24hrs. PBS was added in the surrounding wells to prevent evaporation of media. The media was then aspirated, and 100 µl of the diluted drug with 10 mg/ml doxycycline was added to the corresponding well and incubated for 72 hrs. An MTS assay (CellTiter 96® Promega) was then performed as above to measure the numbers of surviving cells present. Results were analyzed using Graphpad Prism. Non-linear regression was used to calculate IC50 values.

### Radiation

To assess the effect of radiation on our CHRDL2 overexpressing cells, RKO cells were plated at a density of 1x10^5^ in 10 cm round dishes and treated with DMSO control or CHRDL2 ++ doxycycline treatment. After 24 hrs, cells were irradiated using x-ray irradiation at 0 GY, 2 GY, 4 GY and 6 GY. The media and CHRDL2++ treatment was refreshed and cells were incubated under standard conditions for a further 48hrs. Subsequently cell number was counting using Trypan blue cell viability assay (T10282, Thermo Scientific, US).

### Immunofluorescence

For cellular protein detection, cells were plated on coverslips and grown to ∼70% confluency. Cells were fixed with methanol, and the cellular membrane was permeabilized with TRITONX 0.5% for 5 minutes. Cells were blocked with 1% BSA for 1 hr at 37◦C, and then incubated with primary antibodies in 1% BSA for 1hr at 37◦C. A secondary antibody was then added to cells for 1hrs at 37◦C. 5 μl of mounting media with DAPI (VECTASHIELD® WZ-93952-27, Cole-Parmer, UK) was then placed onto the coverslips, and coverslips were fixed on to slides for imaging. Images were obtained using a Leica DM4000 system and corrected total cell fluorescence obtained using ImageJ.

**Antibodies used**:

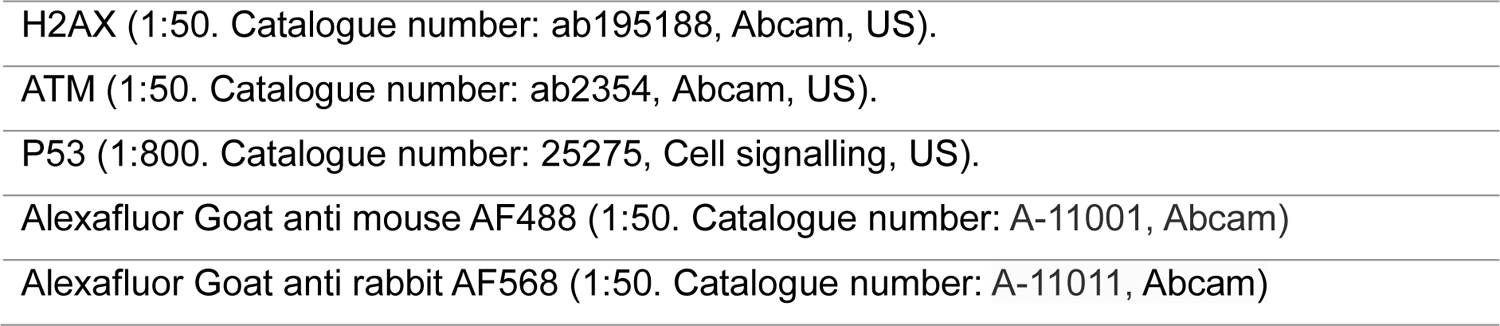

|  |
| --- |
| H2AX (1:50. Catalogue number: ab195188, Abcam, US). |
| ATM (1:50. Catalogue number: ab2354, Abcam, US). |
| P53 (1:800. Catalogue number: 25275, Cell signalling, US). |
| Alexafluor Goat anti mouse AF488 (1:50. Catalogue number: A-11001, Abcam) |
| Alexafluor Goat anti rabbit AF568 (1:50. Catalogue number: A-11011, Abcam) |

### Comet assay

COLO320 cells were plated at a density of 1x10^5^ in with DMSO control or CHRDL2 ++ doxycycline and treated with IC50 oxaliplatin. After 72hrs, cells were harvested and diluted in PBS at a density of 10^4^/ml. Cell suspension was mixed 1:5 with low-gelling agarose and 100 μl was placed on Polylysine coated slides dipped in 1% agarose (Sigma-Aldrich) and allowed to set. A further 100 μl low gelling agarose (Sigma-Aldrich) was placed on top and allowed or set. Cell lysis was then performed by submerging slides in lysis buffer (2% N-Lauroylsarcosine sodium salt (Sigma-Aldrich), 0.5M NA_2_EDTA (Sigma-Aldrich) and 0.1mg/ml proteinase K) for 1hr. Slides were then washed in Electrophoresis buffer for 1.5 hrs (90 mM Tris Buffer (Fisher Bioreagents), 90 mM boric acid (Sigma-Aldrich) and 2 mM NA_2_EDTA). Slides were placed in an electrophoresis tank and submerged in electrophoresis buffer, under 20V current for 40 mins. Slides were then stained with 1% SYBR SAFE (Invitrogen, Thermo Fisher, USA) in TBE for 20 mins, before dehydration through submersion in 70%, 90% and 100% ethanol. Slides were visualised by Leica DM4000 system and tails measured using ImageJ.

### Organoid preparation, culture, and maintenance

Organoids were maintained in an humified atmosphere at 37◦c with 4% CO2. Organoids were grown in ADF media as described: Advanced DMEM/F12, 2mM GLUTAMAX, 1mM N-Acetylcysteine, 10mM HEPES. Supplemented with 1% PS, 10% B27, 5% N2. Growth factors were also given to media surrounding basement membrane extract (Cultrex, Biotechne): 1% Mouse recombinant Noggin (Peprotech, UK), 1% mouse recombinant EGF (Invitrogen, 5% Recombinant human R-spondin (Peprotech).

Organoids were generated as described [34]. Briefly, crypts were isolated from murine small intestine from wild-type mice and washed with PBS. Villi and differentiated cells were scraped off intestine using a glass microscope slide. Sections of intestine were cut into 2mm segments and transferred to ice-cold PBS. Pipettes were coated in FBS, and intestinal segments were washed through pipetting up and down to dislodge single cells and debris. PBS was removed, and washes were repeated 5 times. Segments were then resuspended in 2.5mM EDTA/PBS to loosen crypts and rotated at 4°Cfor 30 mins. The supernatant was then removed, and segments were resuspended in ADF media. The entire volume was pipetted up and down several times, and then the supernatant removed and centrifuged for 5 min at 1200 rpm at 4°C. The supernatant was removed, and the resulting pellet was resuspended in 10ml ADF media and passed through a 70 µm cell strainer into a clean 15 ml falcon tube. The tube was then centrifuged for 2 min at 600 rcf at 4°C so that single cells will not be included in the pellet, and the supernatant was removed. This was repeated 3 times. Finally, the pellet was resuspended in 50ul ADF media and 100ul Cultrex, and pipetted 40ul/ well. Passaging of organoids was repeated every 48hr and consisted of transferring organoid to a 15 ml conical tube, pipetting up and down to break up organoids. Organoids were then centrifuged for 2 min at 600 - 800 rpm at 4°C, and then resuspended in ADF with Cultrex as described previously.

### Organoid immunofluorescence staining

Organoid samples were prepared for staining by removal of growth media and pelleted through centrifugation at 600g. Organoids were then fixed through resuspension in 500 μl neutral buffered formalin (Sigma-Aldrich) for 10 minutes, before pelleting at 400g and resuspension in 70% ethanol for 1 minute. Organoids were then pelleted at 400g and resuspended in 50 μl of low gelling agarose (Sigma-Aldrich) and incubated on ice for 30 minutes, before embedding in paraffin blocks using standard protocols. Sectioning of organoids was performed at 5 μM through standard microtome sectioning and left to dry on slides.

Slides containing organoid sections were dewaxed through xylene (Fisher Bioreagents) submersion for 5 minutes and rehydrated through submersion in ethanol at 100% 90% and 70% for 5 minutes. Antigen retrieval was performed by submerging slides in boiling 10 mM sodium citrate buffer (Sigma-Aldrich), before washing with PBS. Samples were then blocked though addition of Goat serum (Zytochem Plus, 2bscientific, UK) for 1 hr. Primary antibodies diluted in PBS were added for 1hr, and secondary antibodies were incubated for 1hr in the dark. Coverslips were mounted using VECTASHIELD Vibrance [TM] Antifade Mounting Medium with DAPI (2bscientific) for imaging. Slides were visualised by Leica DM4000 system. Organoid staining was scored on a scale of 1-5 by an independent blinded researcher.

**Antibodies used**:

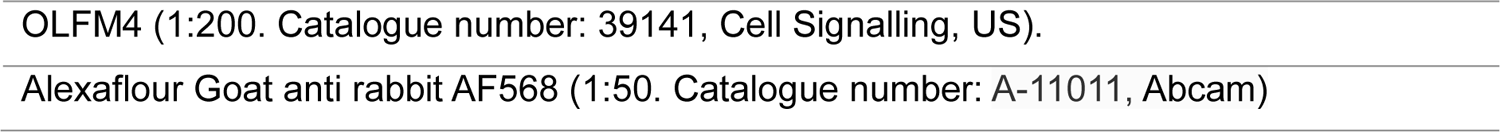

### RNAseq

Samples for RNA-seq analysis were prepared by culturing cells in standard media conditions with overexpression of the CHRDL2 gene through doxycycline-inducible expression. Doxycycline was given in quantities of 10 µg/ml, 1 µg /ml, and 0.1 µg /ml or DMSO control as described previously.

RNA-seq was performed by the Oxford Genomic centre. Data for bioinformatics analysis was given in the format of fastq raw reads. Data was analysed using the open-source software package Tuxedo Suite. Tophat2 and Bowtie2 were used to map paired end reads to the reference Homo sapiens genome build GRCh38. GENCODE38 was used as the reference human genome annotation.

Aligned reads were filtered for quality using Samtools with the minimum selection threshold of 30. Transcripts assembly and quantification was done through Cufflinks, and differential expression analysis was achieved through the use of Cuffdiff software. Differential expression was expressed in the form of log2 fold change between sample and control.

### Data visualisation and R

Data was cleaned and significant data was extracted using R software. Graphs we generated using R studio 4.1.0 using libraries ggplot2 and heatmap2.

Gene-set-enrichment analysis was performed using the GSEA software 4.2.3. The Chip annotation platform used was Human_Ensembl_Transcript_ID_MSigDB.v7.5.1.chip.

Gene sets used:

- c6.all.v7.5.1.symbols.gmt
- h.all.v7.5.1.symbols.gmt
- GOBP_REGULATION_OF_BMP_SIGNALING_PATHWAY
- enplot_REACTOME_PI3K_AKT_SIGNALING_IN_CANCER_13
- enplot_GOMF_BMP_RECEPTOR_BINDING_58
- WP_NRF2_PATHWAY.v2023.1.Hs.

## Acknowledgments

We are grateful to Hayley Davis for her help with establishing intestinal organoids. We thank Ian Tomlinson and Cristina Pina for their advice and comments on the study design and manuscript and Cristina Pina for help with flow cytometry analysis. Thanks also to Jasmin Sandhu for cloning the overexpression constructs for CHRLD2 and Emine Efendi for immunofluorescence scoring.

## Abbreviations

CRC: Colorectal cancer
CHRDL2: Chordin-like 2
BMP: Bone-morphogenic protein
WNT: WNT signalling pathway
ISC: Intestinal stem cell
CSC: Cancer stem cell
LGR5: Leucine-rich repeat-containing G-protein coupled receptor 5
LGR6: Leucine-rich repeat-containing G-protein coupled receptor 6
BMI1: B lymphoma Mo-MLV insertion region 1 homolog)
DSB: Double stranded Break
ATM: Ataxia-telangiectasia mutated
γH2AX: H2A histone family member X

## Author Contributions

A.L. and E.C conceived the study. E.C carried out cell line and organoid studies, and carried out RNAseq analysis. A.L. provided resources and expertise. E.C. and A.L. wrote the manuscript.

