## Supplementary information for "BMP antagonist CHRDL2 enhances the cancer stem-cell phenotype and increases chemotherapy resistance in Colorectal Cancer"

### Supplementary 1: P-SMAD1/5 protein quantification

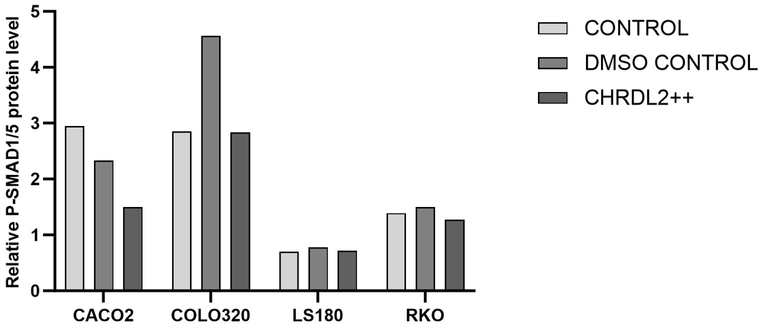


Supplementary figure 1: Quantification of P-SMAD1/5 protein levels in CRC cells with CHRDL2 overexpression.

### Supplementary 2: Quantification of Clonogenic, migration and cell cycle analysis assays

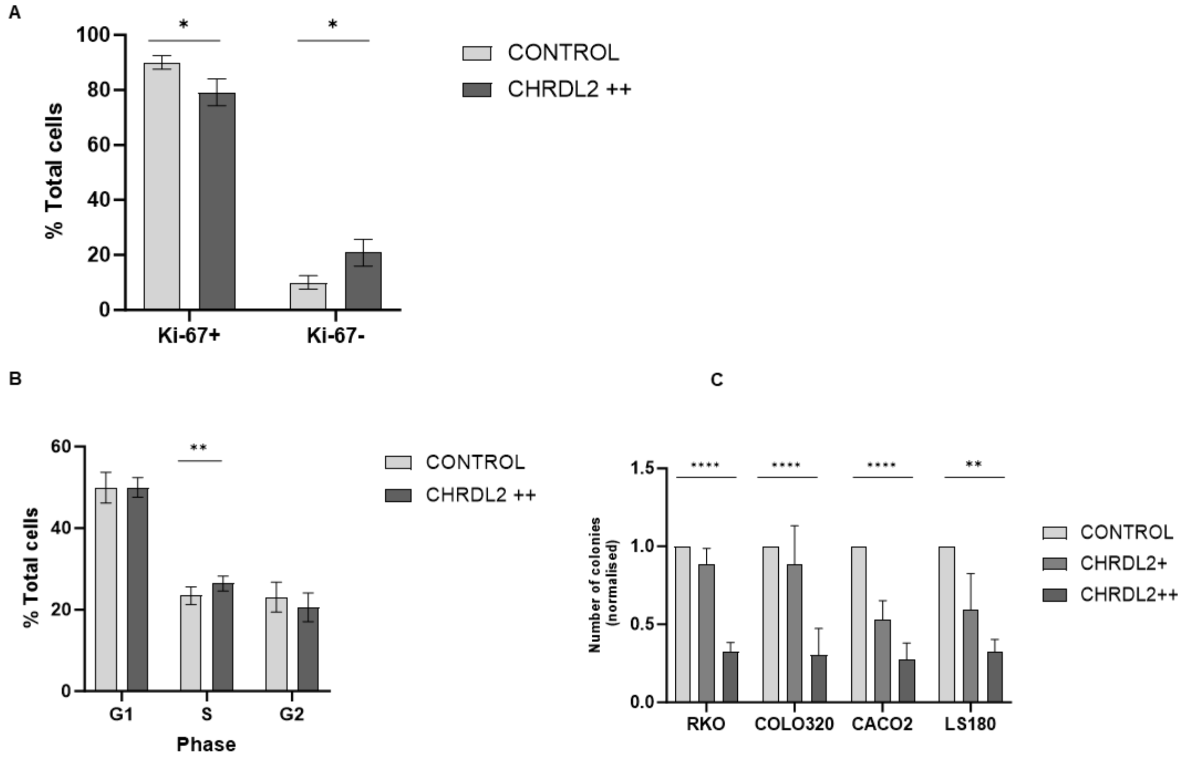


Supplementary figure 2: .A) Quantification of Ki67+/- cells by flow cytometry in COLO320 control and CHRDL2++ cells, P<0.0485. B) Quantification of cell cycle status by flow cytometry in COLO320 control and CHRDL2++ cells, p=0.0076. C) Quantification of clonogenic colonies established in our 4 experimental cell lines with CHRDL2 overexpression. CACO2 and RKO cell lines both showed reduced colony formation in the low and high CHRDL2 treated groups, RKO p<0.01,COLO320 p<0.001, CACO2 p<0.05, LS180 P<0.001, T-test. COLO320 and LS180 both showed a reduction in colony formation in the high CHRDL2 group only, p<0.01. N=3

### Supplementary 3: Cell line IC50s

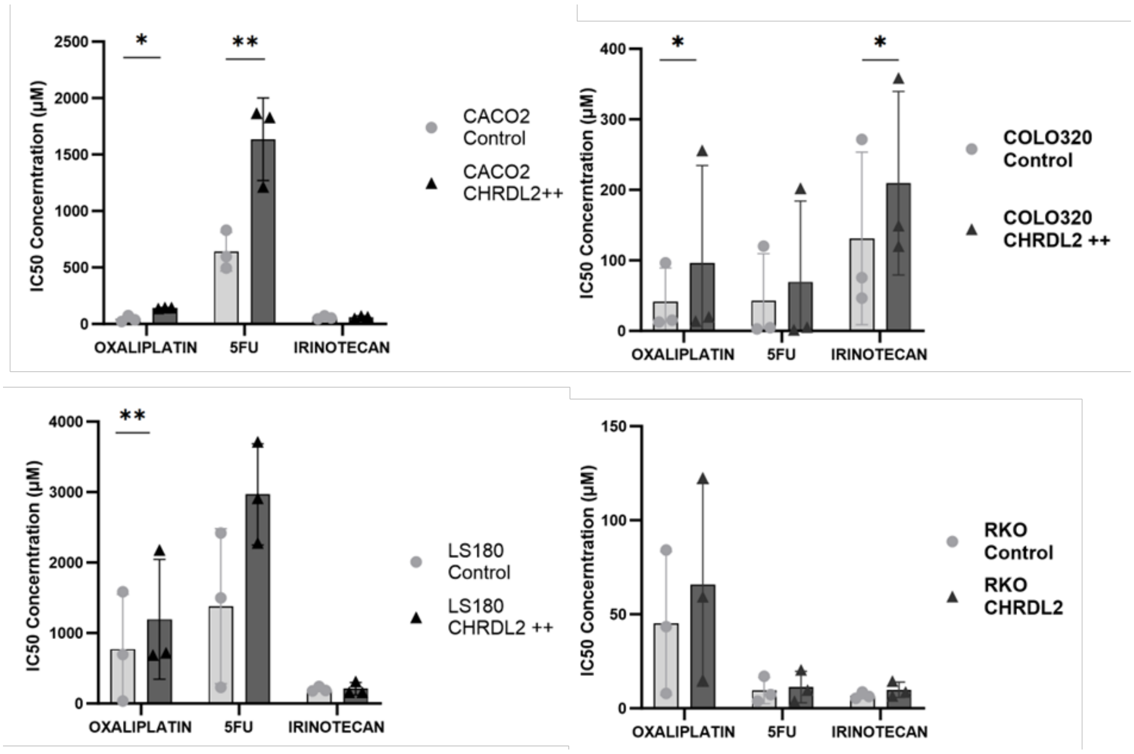


Supplementary figure 3: Average IC50 values of CACO2, COLO320, LS180 and RKO cell lines using chemotherapy drugs Oxaliplatin, 5FU, and Irinotecan N=3.

**Supplementary 4: Ki67 immuno-fluorescence and flow analysis of COLO320 cells treated with chemotherapy Oxaliplatin and Ku70 analysis.**

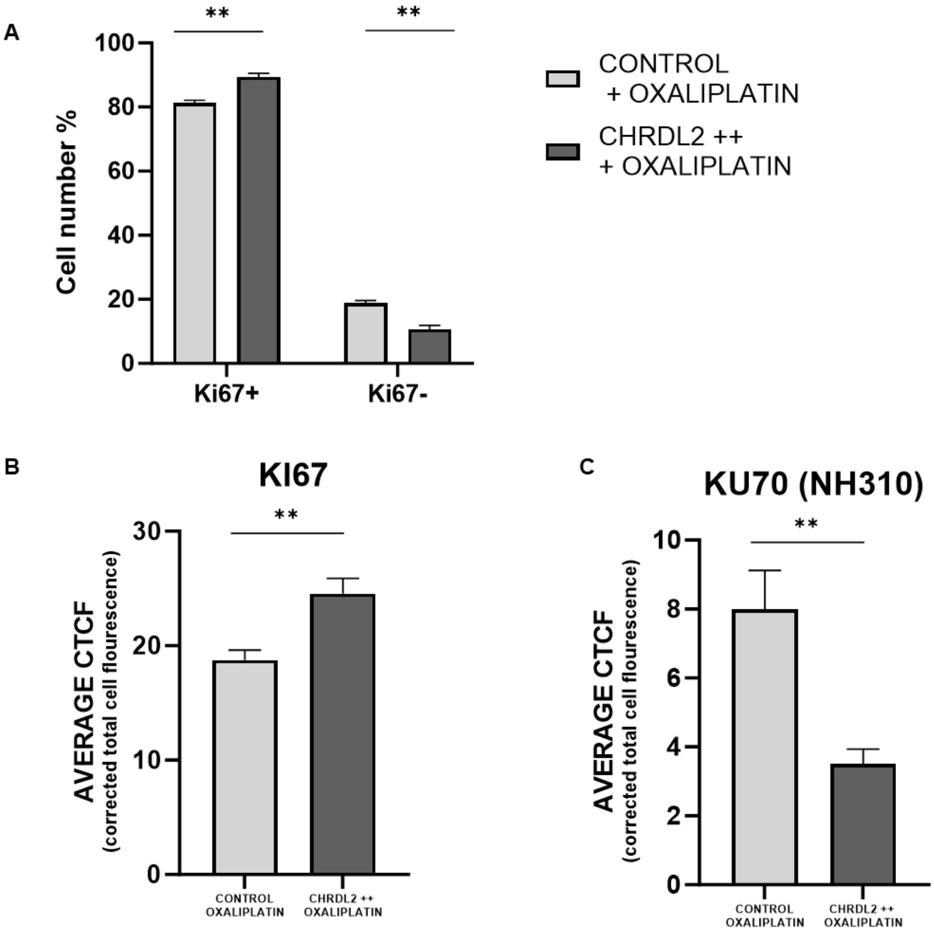


Supplementary figure 4: A) Flow cytometry analysis of Ki67 expression by COLO320 cells treated with 5 μM oxaliplatin.t-TEST. P<0.0056, p<0.0055. n=3 B) Quantification of Ki67 in COLO320 cells overexpressing CHRDL2 treated with 5 μM oxaliplatin 72 hrs. Cells were treated with DMSO control reagent, or Doxycycline to induce CHRDL2 overexpression. P<0.0064. C) Quantification of Ku70 in COLO320 cells overexpressing CHRDL2 treated with 5 μM oxaliplatin 72 hrs. Cells were treated with DMSO control reagent, or Doxycycline to induce CHRDL2 overexpression. P<0.0057 N=3.

**Supplementary 5: P53 Immunofluorescence**

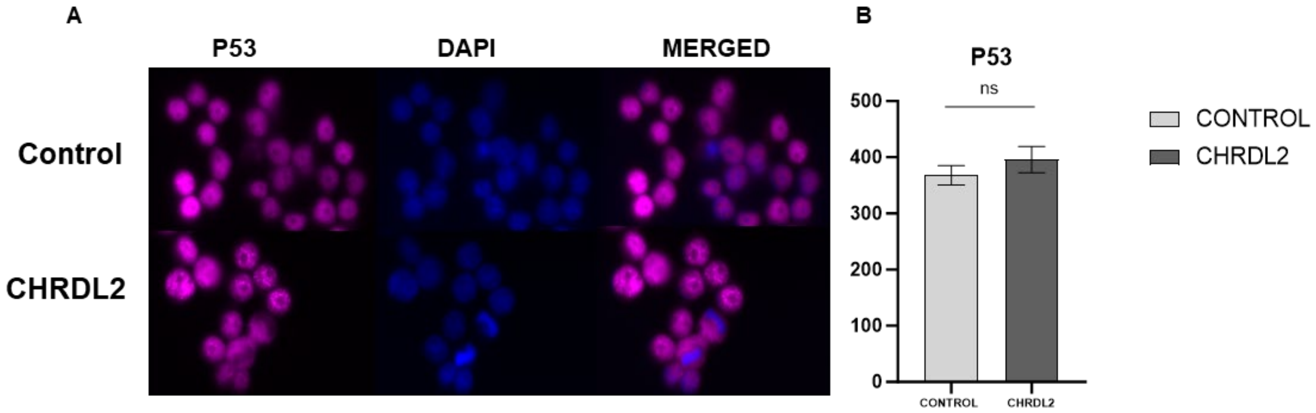


Supplementary figure 5: A) Immunofluorescence staining of P53 on COLO320 cells treated with 5 μM oxaliplatin at 72 hrs. B) Quantification of P53 in COLO320 cells overexpressing CHRDL2 treated with 5 μM oxaliplatin 72 hrs. Cells were treated with DMSO control reagent, or Doxycycline to induce CHRDL2 overexpression. N=3.

### Supplementary 6: GSEA plots

#### WNT Signalling

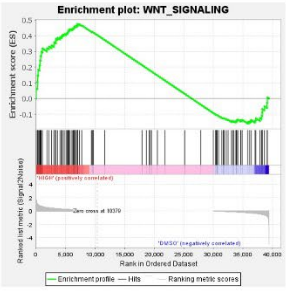


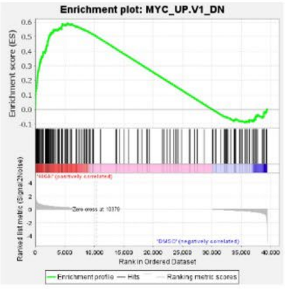


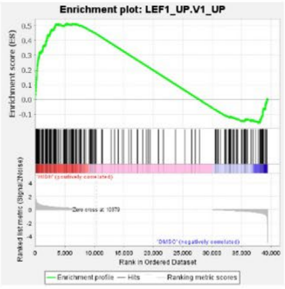


#### YAP Signalling

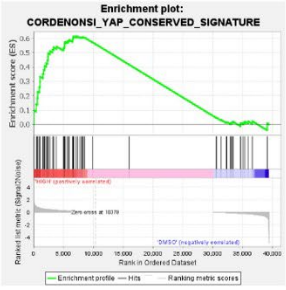


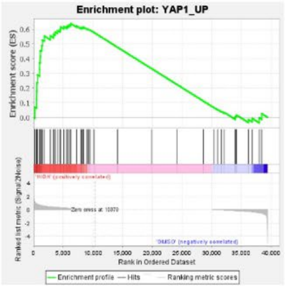


#### BMI1 Signalling

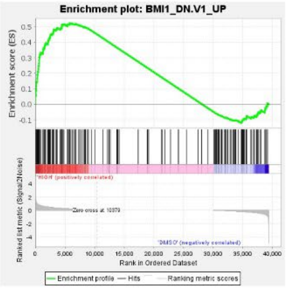


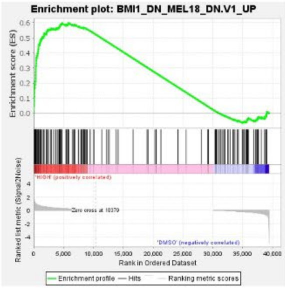


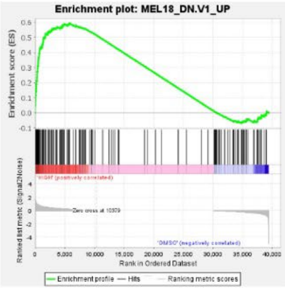


#### RAF Signalling

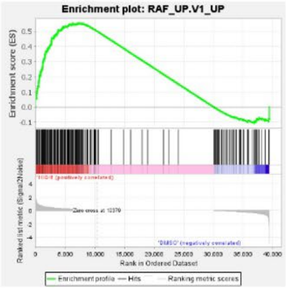


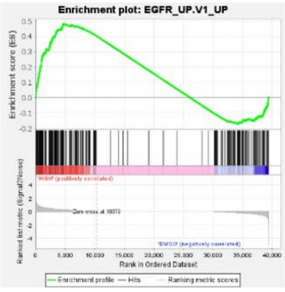


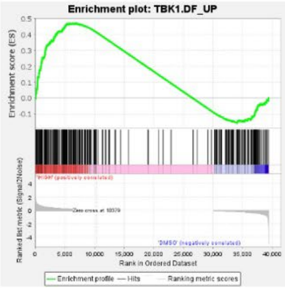

### Metabolism

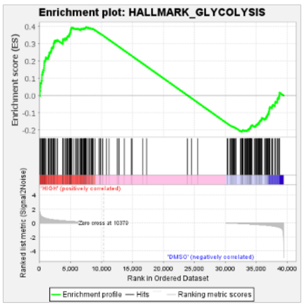


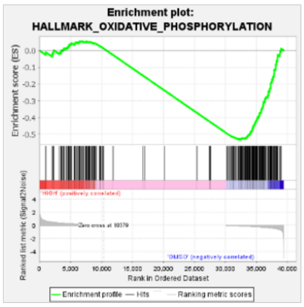


### DNA damage repair

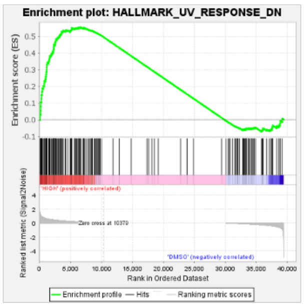


### Cell cycle

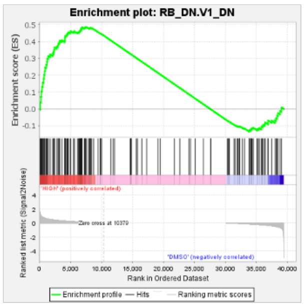


### MTOR signalling

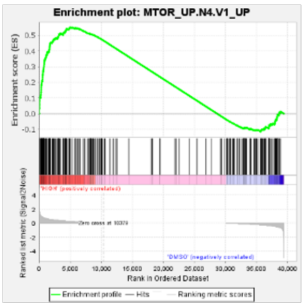


### Hallmark cancer pathways

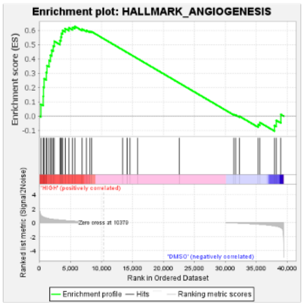


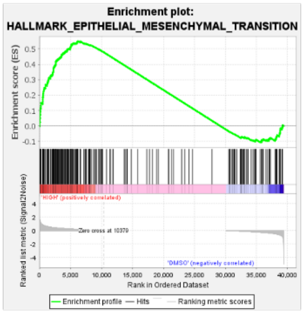


### cAMP signalling

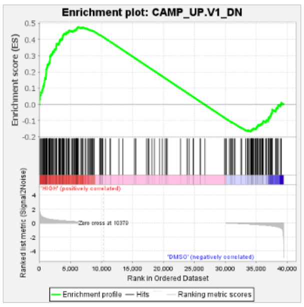


### ALK signalling

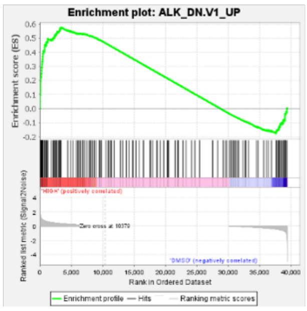


### IL2/2TAT5 signalling

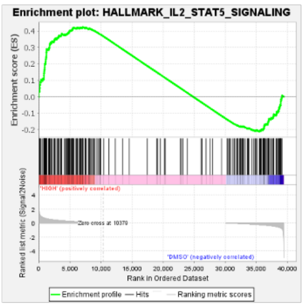


Supplementary figure 6: GSEA plots of RNAseq analysis from CHRDL2++ cells (HIGH) compared to DMSO control. P values given as FWER. WNT_SIGNALLING NES= 1.15 P<0.0. MYC_UP.V1_DN NES=1.25 P<0.01. LEF1_UP_V1_UP NES=1.26 P<0.01. CORDENOSI_YAP_CONSERVED_SIGNATURE NES=1.36 P<0.01. YAP1_UP NES=1.32 P<0.01. BMI1_DN.V1_UP NES=1.33 P<0.01. BMI1_DN_MEL18_DN.V1_UP NES=1.38 P<0.01. MEL18_DN.V1_UP NES=1.41 P<0.0. RAF_UP.V1_UP NES=1.2 P<0.01. EGFR_UP.V1_UP NES= 1.25 P<0.0. TBK1.DF_UP NES=1.2 P<0.01. HALLMARK_GLYCOLYSIS NES=1,13 P<0.01. HALLMARK_OXIDATIVE_PHOSPHORYLATION NES=-1.1 P<0.01. HALLMARK_UV_RESPONSE_DN NES=1.199 P<0.01. RB_DN.V1_DN NES=1.2 P<0.01. MTOR_UP.N4.V1_UP NES=1.31 P<0.01. HALLMARK_ANGIOGENESIS NES=1.21 P<0.01. HALLMARK_MESENCHYMAL_TRANSITION NES=1.29 P<0.01. CAMP_UP.V1_DN NES=1.18 P<0.01. ALK_DN.V1_UP NES=1.36 P<0.01. HALLMARK_IL2_STAT5_SIGNALLING NES=1.15 P<0.01. N=3.
